# Trajectory Inference for Multi-Omics Data Using Ordered Labels

**DOI:** 10.1101/2025.02.25.640243

**Authors:** Hiroshi Kobayashi, Masaaki Okabe, Hiroshi Yadohisa

**Affiliations:** Graduate School of Culture and Information Science, Doshisha University, Japan; Faculty of Culture and Information Science, Doshisha University, Japan

**Keywords:** Pseudotime estimation, Data integration, Variational Autoencoder

## Abstract

Trajectory inference (TI) has emerged as a crucial approach for understanding cellular development and differentiation processes. By reconstructing the temporal ordering of individual cells, TI can analyze the mechanisms underlying development and disease progression. Various TI methods have been proposed and applied across diverse fields, including cancer research and immunology. However, most existing methods rely solely on gene expression data and do not incorporate other omics information such as DNA methylation, chromatin accessibility, and protein expression. To address this limitation, approaches that integrate multiple omics datasets for TI have been proposed. Nevertheless, these methods typically assume the use of only common variables across datasets, making it difficult to align heterogeneous data with differing measurement targets across omics layers. This limitation can lead to information loss and challenges in capturing interactions between different omics datasets, which may reduce integration accuracy. In this study, we propose a novel TI method that captures interactions among variables across different omics layers and accommodates unpaired datasets with non-overlapping measurement targets. By effectively capturing relationships between different omics layers, our approach enables a more comprehensive multi-omics TI framework, thus providing deeper insights into cellular development.

## 1 Background

Trajectory inference (TI) is a crucial approach for understanding cellular differentiation and developmental processes [1, 2, 3]. TI reconstructs the temporal trajectory of individual cells to model the processes of development and differentiation. To date, various TI methods have been developed, such as Monocle3, Slingshot, and PAGA. These methods can infer cellular trajectories based on single-cell gene expression data, thereby revealing differentiation pathways. In contrast, supervised approaches, such as psupertime [4], utilize predefined temporal ordering. psupertime improves trajectory estimation accuracy by leveraging prior temporal knowledge and gene expression data. These TI methods have contributed significantly to advancing our understanding of cellular development and disease progression, providing novel insights across various fields, including cancer research and immunology [1, 3]. However, most existing TI methods rely on only single-cell RNA sequencing (scRNA-seq) data and do not incorporate other omics information [5]. For instance, changes in DNA methylation, chromatin accessibility, and protein expression significantly influence cellular states, yet conventional single-omics TI methods cannot account for these regulatory influences. Therefore, to gain a more comprehensive understanding of cellular dynamics, it is essential to integrate multiple types of omics data rather than rely solely on gene expression profiles. The integration of multiple omics datasets is referred to as multiomics data [6]. Multi-omics analysis provides a more comprehensive approach to understanding complex biological systems and diseases compared to single-omics analysis [7].

Considering these challenges, VITAE [8] has been introduced as a TI framework to integrate multi-omics data. VITAE integrates multiple omics modalities, such as scRNA-seq and single-cell assay for transposase-accessible chromatin sequencing (scATAC-seq), in the TI process. However, VITAE requires preprocessing to restrict the analysis to shared variables across datasets, limiting its applicability to datasets with consistent measurement variables across omics layers. For instance, if measurement targets are not aligned across omics layers, variable matching becomes necessary, which can cause information loss. Additionally, this constraint hinders the comprehensive modeling of interactions between different omics layers. Thus, to enable effective TI with multi-omics data, more robust embedding strategies are needed to integrate diverse types of omics information. Several approaches have integrated multi-omics data, including Joint and Individual Variation Explained (JIVE) [9] and Multi-Omics Factor Analysis (MOFA) [10]. JIVE integrates multiple omics datasets by separating shared and individual components. MOFA employs a probabilistic latent variable model to learn common latent structures among various omics datasets to achieve integrated analysis. However, these methods face challenges when omics datasets have misaligned measurement targets (rows) or feature sets (columns). Conventional integration methods are difficult to apply to unpaired datasets with heterogeneous measurement targets and feature definitions across omics layers. Such limitations may impair the ability of TI to effectively capture cross-omics interactions.

Therefore, this study aims to develop a novel TI method that accounts for inter-variable interactions in multi-omics data and accommodates unpaired datasets with misaligned rows and columns. We propose a novel pseudotime estimation algorithm based on Graph-Linked Unified Embedding (GLUE) [11], called CGLUE-SOE. As shown in Table 1, GLUE is capable of handling unpaired datasets, distinguishing it from other methods. Furthermore, GLUE has been empirically demonstrated in the benchmark study by [12] to outperform other methods, achieving superior integration accuracy. We extend GLUE with the motivation of data integration with Ordered Labels for TI applications. Specifically, we leverage GLUE to capture interactions among omics variables while introducing a new objective function and incorporating ordered labels as constraints to derive a trajectory-aware low-dimensional representation. This approach enables TI in datasets with misaligned measurement targets and variables, thereby enhancing understanding of cellular development.

**Table 1:** Comparison of different methods.

| Method | Embedding | Integration | Unpaired | TI | Ordered Label |
| --- | --- | --- | --- | --- | --- |
| CGLUE-SOE (Proposed) | ✓ | ✓ | ✓ | ✓ | ✓ |
| GLUE | ✓ | ✓ | ✓ |  |  |
| JIVE | ✓ | ✓ |  |  |  |
| MOFA | ✓ | ✓ |  |  |  |
| VITAE | ✓ | ✓ |  | ✓ |  |
| CCA | ✓ | ✓ |  |  |  |

Figure 1 presents a conceptual overview of the proposed method. This model accepts datasets with misaligned rows and columns as input, along with ordered labels assigned to the targets. It maps all targets onto a shared lowdimensional embedding space, thereby enabling two-dimensional visualization and pseudotime estimation. Figure 2 (B) and (C) illustrate the differences in embeddings produced by existing methods and our proposed method. The objective of our proposed method is to generate embeddings that resemble those depicted in the right panel of Figure 2 (B) and (C). In Figure 2 (B), the left panel represents the embedding results of existing methods, where data points are projected into a low-dimensional space without considering label information, resulting in widely dispersed embeddings of samples sharing the same label. In contrast, the right panel shows the results from our proposed method, where label information is incorporated to encourage well-clustered embeddings of samples with the same label. By incorporating label information into the embedding process, this approach yields more interpretable embeddings that facilitate downstream analyses, such as clustering and classification. The left panel of Figure 2 (C) illustrates another limitation of existing methods. Even if data points are grouped according to their labels, failing to account for the sequential relationship among labels may result in embeddings that contradict the inherent order. For instance, in developmental or differentiation processes, cell populations expected to follow a sequential order may be embedded inconsistently. In contrast, the right panel shows the results of our proposed method, where the loss function is specifically designed to preserve sequential relationships in the embedding space, ensuring that the order among labels is maintained. Thus, the proposed method incorporates label information and preserves order structure, resulting in more interpretable embeddings. This study presents a comprehensive description of our method and evaluate its effectiveness through experiments on both simulated and real-world datasets.

**Figure 1:**
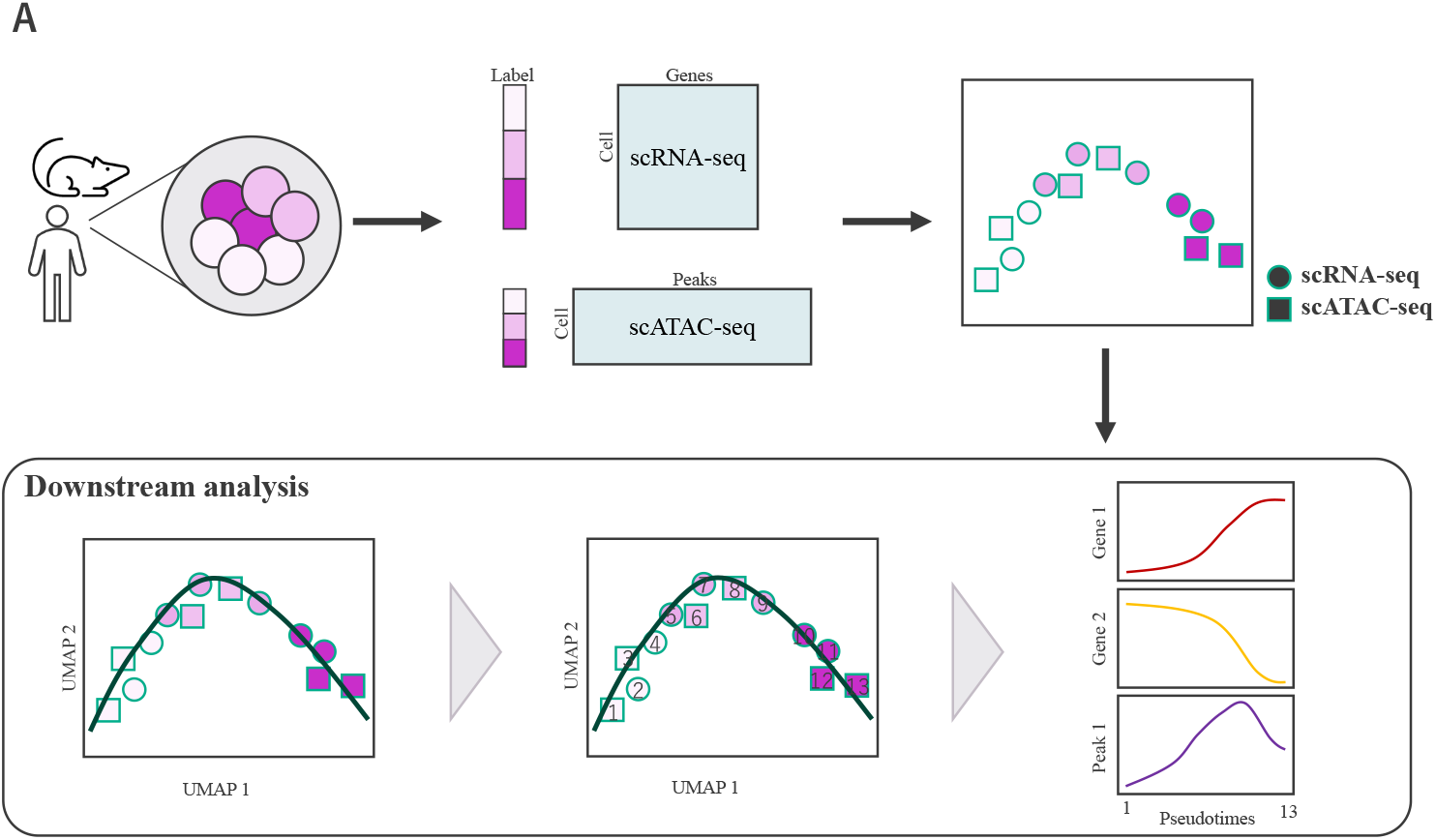
(A) Overall structure of the model. Using multi-labeled omics data as input data, our method can obtains a low-dimensional structure awaring the information of the labels. The obtained low-dimensional structure enables downstream analysis such as trajectory inference and pseudotime estimation.

**Figure 2:**
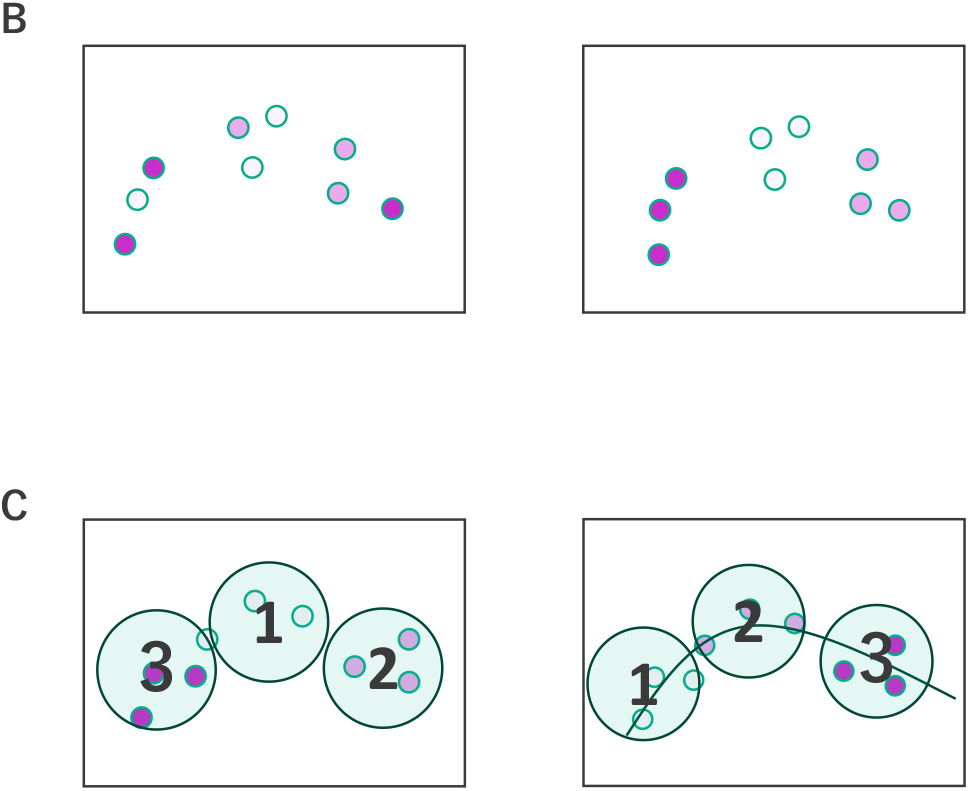
(B) The first issue with existing methods: The left panel shows an example of incorrect embedding, while the right panel represents the ideal embedding, which aims for well-clustered embeddings per label. (C) The second issue with existing methods is shown in the left panel, which presents incorrect embedding, while the right panel displays the ideal embedding, which aims to preserve the correct order structure.

## 2 Discussion

Conditional Graph-Linked Unified Embedding with Soft Ordinal Embedding (CGLUE-SOE) incorporates two key innovations for integrating multi-omics data while leveraging available ordered labels. The first leverages label information assigned to each sample as a supervisory signal during embedding to allow samples sharing the same label to be embedded in close proximity in low-dimensional space. This approach yields embeddings that explicitly encode label information, thereby facilitating label structure-based interpretation. The second entails computing centroids in the low-dimensional space for each ordered class label, such as disease stages or cell maturation levels, following embedding. A triplet loss term is introduced to penalize cases where the distance relationships between these centroids contradict the expected order of class labels. By incorporating this constraint, the learned embeddings maintain sequential relationships among labels, allowing for the interpretation of ordered structures in dynamic biological processes such as cell differentiation and development. A more detailed explanation of these mechanisms is provided in Chapter 4.

### 2.1 Real datasets

For real data applications, we have utilized a dataset published by [12]. This dataset contains five distinct cell types with ordered labels that primarily represent the differentiation trajectory of intermediate progenitor cells to upper-layer excitatory neurons Table 2 summarizes the results of evaluating each method on the real dataset across 10 independent runs. Among all methods, CGLUE-SOE achieved the highest accuracy, as measured by Spearman’s rank correlation coefficient (Correlation) and Adjusted Rand Index (ARI) scores. This was followed by Conditional Graph-Linked Unified Embedding (CGLUE), GLUE-SOE, and Graph-Linked Unified Embedding (GLUE). Regarding the standard deviation of Correlation, both CGLUE-SOE and GLUE-SOE had the lowest values. For the standard deviation of ARI, GLUE-SOE, and GLUE exhibited the least variability.

**Table 2:** Mean and standard deviation (Sd) of accuracy metrics on the real dataset.

|  | Correlation |  | ARI |  |
| --- | --- | --- | --- | --- |
|  | Mean | Sd | Mean | Sd |
| CGLUE-SOE | **0.644** | 0.012 | **0.250** | 0.036 |
| GLUE-SOE | 0.619 | 0.012 | 0.242 | 0.010 |
| CGLUE | 0.633 | 0.021 | 0.249 | 0.033 |
| GLUE | 0.608 | 0.018 | 0.239 | 0.014 |

Then, we present the embedding results with the highest accuracy for both the proposed (CGLUE-SOE) and existing methods (GLUE) in Figures 3a and 3b, respectively. The ordered relationship among the labels is defined as ≺ IP_Hmgn2 IP_Gadd45g ≺ IP_Eomes ≺ Ex23_Cntn2 ≺ Ex23_Cux1. The black curve in the figures represents the principal trajectory estimated by Slingshot, which defines a pseudo-temporal axis, and the cross markers denote the centroid positions of each label. Based on the visualization in this two-dimensional space, CGLUE-SOE achieves a clearer separation of labels.

**Figure 3:**
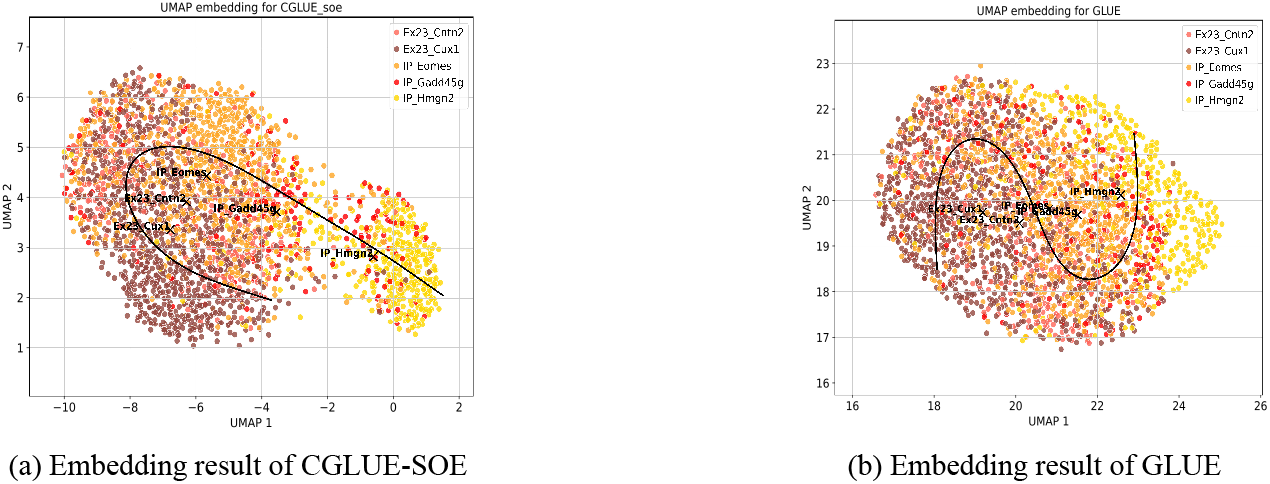
Comparison of embedding results between CGLUE-SOE and GLUE

Next, we estimated the pseudotime for each sample based on the embedding results and principal trajectory estimated by Slingshot to construct histograms (Figures 4 and 5), with the x-axis representing pseudotime and y-axis indicating the number of samples assigned to each pseudotime.

**Figure 4:**
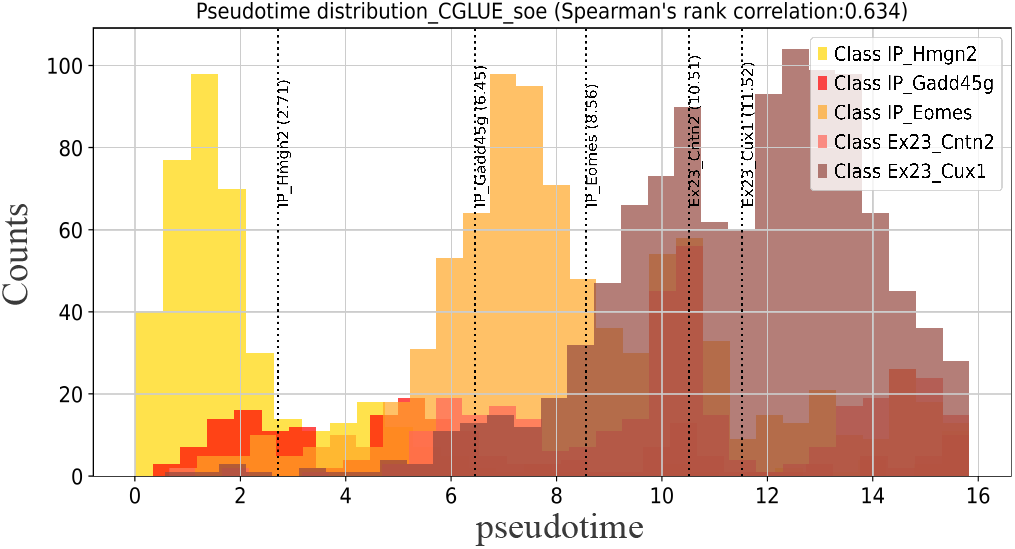
Pseudotime estimation results of CGLUE-SOE

**Figure 5:**
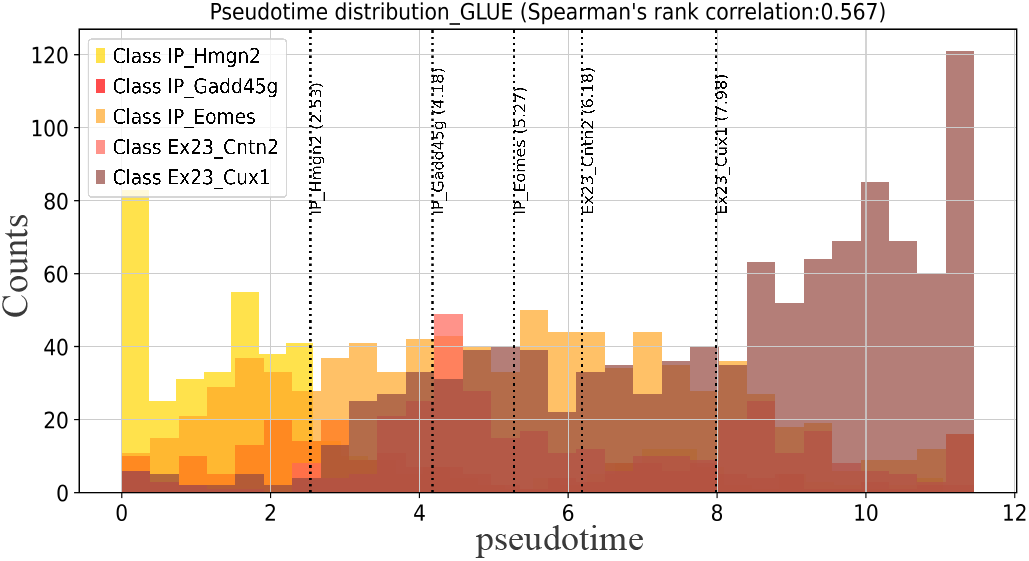
Pseudotime estimation results of GLUE

**Figure 6:**
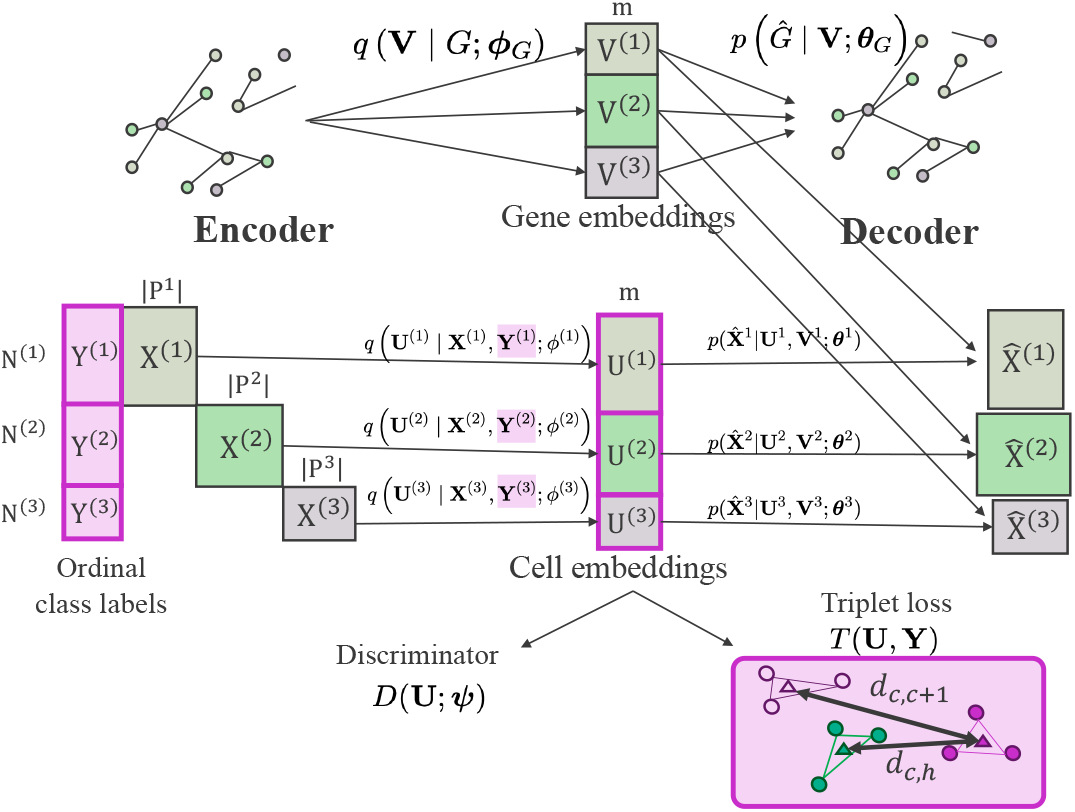
Model architecture. The input consists of three omics datasets, 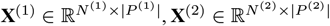, and 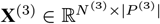, along with categorical labels for each sample, encoded as one-hot vectors: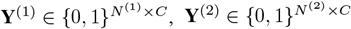, and 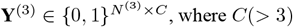, where *C*(*>* 3) represents the number of label categories. Here, *N*^(1)^, *N*^(2)^, and *N*^(3)^ denote the number of samples (cells), and |*P*^(1)^|, |*P*^(2)^|, and |*P*^(3)^| represent the number of features in each dataset. The proposed method embeds each dataset into a shared low-dimensional space. Although the original datasets may have different feature dimensions and distributions, the embedding dimension, denoted as *m*, is shared across all datasets. Relationships between features across different datasets are incorporated using prior knowledge in the form of a guidance graph *G* = (*P, E*), which encodes known interactions. Then, the decoder reconstructs the original omics data by computing the inner product between cell and gene embeddings, thereby transforming the omics-specific data space into a common space guided by the knowledge graph. To harmonize cell embeddings across different datasets, an adversarial discriminator *D* is introduced to align the embedded representations through adversarial learning. The trainable parameters of the data and graph encoders are denoted as *ϕ*_1_, *ϕ*_2_, *ϕ*_3_, and *ϕ*_*G*_, while those of the data and graph decoders are represented as *θ*_1_, *θ*_2_, *θ*_3_, and *θ*_*G*_. Additionally, *ψ* represents the trainable parameters of the discriminator. Furthermore, triplet loss is incorporated to ensure that the embedding preserves the sequential relationships among labels by leveraging both cell embeddings and ordered label information.

These results revealed well-separated distributions for each label, suggesting that reasonable pseudotime assignments were achieved for each sample and in datasets with ordered labels, CGLUE-SOE can effectively reconstruct the sequential progression of cellular maturation in a low-dimensional space

### 2.2 Simulation study

When describing the overall trends based on data characteristics, we observed that accuracy improved as the number of samples and features increased. Additionally, datasets with balanced label distributions achieved higher accuracy than those with imbalanced ones, and a greater number of label-associated variables was associated with improved accuracy.

Next, we evaluated the performance of the proposed and baseline methods. For datasets with 200 rows, 200 columns, and 5 labels, CGLUE-SOE, the proposed method, achieved the highest Correlation score. Following this, CGLUE-SOE, CGLUE, GLUE-SOE, and GLUE demonstrated high accuracy, and were ranked in a descending order. Contrastingly, in terms of ARI, CGLUE outperformed other methods except when the number of label-associated variables was high, followed by CGLUE-SOE, GLUE-SOE, and GLUE.

For datasets with 500 rows, 500 columns, and 5 labels, with a low number of label-associated variables, CGLUE-SOE demonstrated the highest accuracy both in terms of Correlation and ARI, followed by CGLUE, GLUE-SOE, and GLUE. In contrast, when the number of label-associated variables was moderate, CGLUE-SOE and CGLUE showed the highest accuracy in terms of Correlation, whereas CGLUE had the highest ARI score. These results suggest that, when there are few label-associated variables and the dataset lacks sufficient information, CGLUE-SOE mitigates this limitation by leveraging an order-aware loss term.

Thus, CGLUE-SOE is preferable for small-data matrices or imbalanced label distributions due to its ability to preserve order information. In contrast, when the dataset inherently captures distinctions between labels or prioritizes label-specific clustering accuracy, CGLUE is a more suitable choice. These findings suggest that CGLUE is preferable when the primary concern is clustering accuracy at the label level.

## 3 Conclusions

In this study, we proposed a novel method to address the challenges associated with embedding multi-omics data. The proposed method facilitates label-driven data interpretation by incorporating label information assigned to each sample into the embedding process. Furthermore, it aims to preserve order information when the sequential nature of labels is biologically meaningful, which is particularly important for downstream analyses, such as pseudotime estimation. Specifically, our method incorporates label information as a conditioning factor during embedding, ensuring that samples with the same label are positioned close together in the low-dimensional space. Additionally, by introducing a triplet loss function, our approach faithfully preserves sequential relationships in the low-dimensional space, thus facilitating interpretation of dynamic processes, including disease progression and cell differentiation.

To assess the effectiveness of our method, we conducted numerical experiments and applied this method to real-world datasets. The numerical experiments demonstrated that the proposed method outperforms existing approaches in terms of clustering accuracy and rank correlation, a metric that quantifies the preservation of order information. Analysis using real-world data showed that our method effectively captured label and order information into the embedding of multi-omics data, facilitating the extraction of biologically meaningful insights.

Two major challenges have been noted for future research. First, is addressing cases where only a subset of samples has assigned labels. In this study, we assumed that all samples had assigned class labels. However, in real-world datasets, labels may be missing for some samples, highlighting the need for a conditional embedding approach that can handle missing labels. Addressing this issue would allow our method to be applied to more diverse datasets, further increasing its practical utility. Second, is explicitly modeling order relationships with branching structures, as the proposed method depends on a single linear ordering of labels. However, in biological processes such as cell differentiation and disease progression, branching-order structures frequently occur. Our method does not explicitly account for such branching relationships, potentially limiting its capacity to model complex biological dynamics with multiple differentiation pathways. Developing an embedding approach that captures branching structures would enhance the interpretability of dynamic processes involving multiple differentiation pathways. By overcoming these challenges, we expect this proposed method to have broad applications, facilitating deeper biological insights than those yielded by existing methods.

## 4 Method

### 4.1 Notation

We consider a multi-omics dataset comprising *K* different omics data matrices. The *k*-th omics data matrix is denoted as 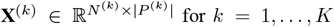 where multiple data matrices are given. Here, the index for samples (e.g., cells) in the *k*-th data matrix is denoted as *n*^(*k*)^ (= 1, …, *N*^(*k*)^), and index for features (e.g., genes) is represented as *i ∈ P*^(*k*)^. Considering these data matrices, we define the embedding of samples into an *m*-dimensional space (*m*≤ min_*k*_(*N*^(*k*)^, |*P*^(*k*)^|)). The embedding matrix for samples in the *k*-th data matrix is denoted as 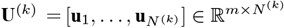, where **u** *∈* R^*m*^ represents the embedding of an individual sample. The embeddings of all samples across all data matrices are concatenated into a single matrix:

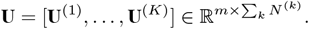

When the same samples are fully observed across different datasets, the integration is referred to as horizontal integration. For instance, considering two omics datasets, 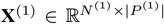 and 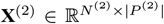, if *N*^(1)^ = *N*^(2)^ and the samples are fully matched, then horizontal integration is performed. Conversely, when differences are observed in either the rows (samples) or columns (features) across datasets, the integration is referred to as diagonal integration, which allows complementary information from different datasets to be leveraged effectively.

We define a knowledge graph *G* = (*P, E*) that consists of the set of all features *P* and a set of edges *E* = *{*(*i, j*) | *i, j ∈P}* representing relationships among features. This knowledge graph encodes prior knowledge regarding known feature interactions (e.g., gene—gene interactions), allowing researchers to incorporate such relationships into the model. Interactions among genes are often well-documented in multi-omics research, thus, our methods considers these interactions. The interaction sign between two features *i* and *j* is represented as *s*_*ij*_ ∈ {−1, 1}, where *s*_*ij*_ = 1 indicates that feature *i* promotes feature *j*, and *s*_*ij*_ = *−*1 indicates an inhibitory effect. The weight of the interaction is denoted as *w*_*ij*_ *∈* (0, 1]. In general, interaction weights are not necessarily symmetric, i.e., *w*_*ij*_ ≠ *w*_*ji*_.

The index for features (e.g., genes) in the *k*-th data matrix is denoted as *i ∈ P*^(*k*)^ and union of all feature sets across datasets is as follows.

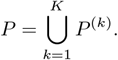

A single sample vector from the *k*-th data matrix is represented as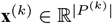. Subsequently, we define the embedding of a feature *i* in the *m*-dimensional space as **v**_*i*_ ∈ ℝ^*m*^. The embedding matrix for features in the *k*-th data matrix is denoted as

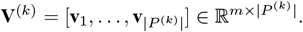

The embeddings of all features across all data matrices are concatenated into a single matrix:

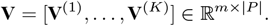

### 4.2 Model

For a given sample, we define its categorical label as a one-hot encoded vector 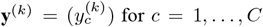 for *c* = 1,..., *C*, where **y**^(*k*)^ *∈* {0, 1}^*C*^ and

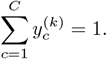

The label vectors for all samples in the *k*-th data matrix are then aggregated into a matrix:

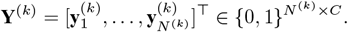

This matrix **Y**^(*k*)^ contains the label information for all samples in the *k*-th dataset. To integrate label information across all datasets, we concatenate the matrices **Y**^(*k*)^ for all *k* = 1, …, *K* into a single matrix:

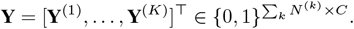

The encoder model is defined as follows.

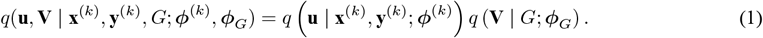

Here, *q* (**u** | **x**^(*k*)^, **y**^(*k*)^; ***ϕ***^(*k*)^) represents the data encoder, while *q* (**V** | *G*; ***ϕ***_*G*_) represents the graph encoder. The vector **y**^(*k*)^ encodes the label information, which is explicitly utilized in our method, unlike existing approaches that do not directly incorporate label or categorical information. The parameters ***ϕ***^(*k*)^ and ***ϕ***_*G*_ correspond to the trainable parameters of the data and graph encoders, respectively.

The graph encoder in Eq. (1) is defined as follows:

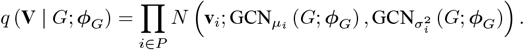

This formulation models the feature embeddings **V** as a multivariate normal distribution whose parameters are learned via a graph convolutional network (GCN). A GCN is a neural network model that propagates information through the graph structure, capturing feature relationships. Here, 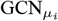 represents the model predicting the mean of the normal distribution whereas 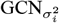 represents the model predicting the variance.

The data encoder in Eq. (1) is defined as follows:

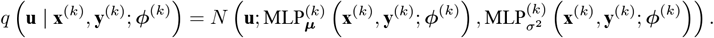

This formulation models the sample embeddings **u** as a multivariate normal distribution whose parameters are learned via a Multilayer Perceptron (MLP). Here, 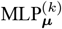 represents the model predicting the mean of the normal distribution, whereas 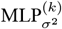 represents the model predicting the variance. By incorporating label information **y**^(*k*)^ alongside input data **x**^(*k*)^, our method generates a label-aware low-dimensional representation, distinguishing it from existing methods that do not explicitly utilize label information.

The decoder model is defined as follows:

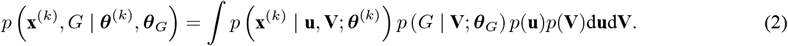

Here, *p* (**x**^(*k*)^ | **u, V**; ***θ***^(*k*)^) represents the data decoder, while *p* (*G* | **V**; ***θ***_*G*_) represents the graph decoder. The parameters ***θ***^(*k*)^ and ***θ***_*G*_ correspond to the trainable parameters of the data and graph decoders, respectively. The term *p*(**u**) denotes the prior distribution of the latent variables for the samples, and *p*(**V**) denotes the prior distribution of the latent variables for the features. These priors are assumed to follow multivariate normal distributions:

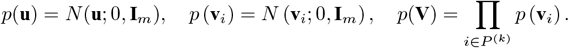

The graph decoder in Eq. (2) is defined as follows:

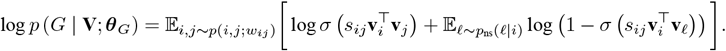

Here, *σ* represents the sigmoid function. The term *p*_ns_ refers to negative sampling, which is used to model non-connected relationships by leveraging features *ℓ* that are indirectly connected to feature *i*. Negative sampling enables contrastive learning by allowing the model to focus on learning connected relationships while learning non-connected relationships. Specifically, edges (*i, j*) are sampled with a probability proportional to their edge weights, and then a feature *ℓ* that is not connected to *i* is sampled for negative contrast.

The data decoder in Eq. (2) varies depending on the assumed data distribution. For instance, to model gene expression counts, a count-based distribution must be assumed. In this case, a negative binomial distribution with dataset-specific parameters is used for each omics dataset, defined as follows:

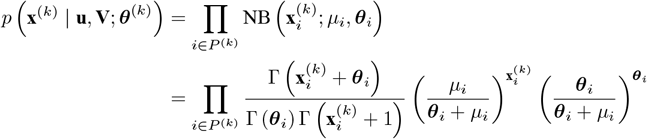

Here, the mean parameter *µ*_*i*_ is defined as:

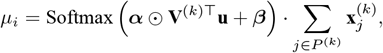

where *⊙* denotes the Hadamard product. The parameters of the negative binomial distribution include 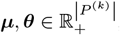, which represent the mean and dispersion parameters, respectively. The scaling coefficient is given by 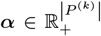, and the bias term is 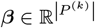. The full set of parameters for the data decoder is denoted as ***θ***^(*k*)^ = *{****θ, α, β****}*.

### 4.3 Objective Function

The objective function consists of four components: the data-related objective function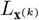, graph-related objective function *L*_*G*_, adversarial learning objective function *L*_D_, and order-preserving objective function *L*_*T*_. The data-related objective function is defined as follows:

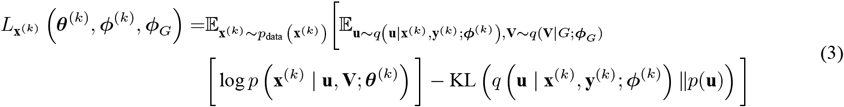

The first term in this objective function represents the negative reconstruction error, corresponding to the log-likelihood of the original data **x**^(*k*)^ considering the latent variables **u** and **V** sampled from the posterior distribution. The second is a regularization term, ensuring that the posterior distribution of **u** remain close to the prior distribution. By maximizing Eq. (3), the latent variable distribution is encouraged to resemble the assumed prior distribution while ensuring that the original data is effectively reconstructed from latent variables. Similarly, the graph-related objective function is defined as:

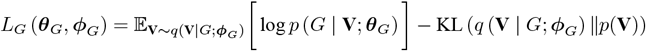

Next, the adversarial learning objective function is defined as follows.

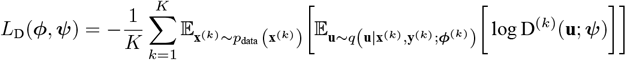

Herein, adversarial learning is instrumental in integrating multiple datasets by enforcing a competition between the discriminator and model. The discriminator attempts to distinguish between different datasets, while the model counteracts this by projecting data from different sources into a shared space. This mechanism ensures that samples obtained from different datasets are embedded into a common low-dimensional space. In CGLUE-SOE, we incorporate the idea from [13] to impose a penalty term that constrains the embedding distances based on the order information among samples. We assume that each sample is assigned one of *C* (≥ 3) ordered labels. The order-preserving objective function is defined as:

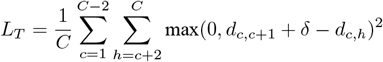

Here, *d*_*c*,*c*+1_ and *d*_*c*,*h*_ represent the Euclidean distances between centroids in the embedding space for order labels *c* and *c* + 1 and for labels *c* and *h*, respectively. The hyperparameter *δ* (*>* 0) is introduced to ensure that the distance *d*_*c*,*c*+1_ between adjacent labels remain sufficiently smaller than the distance *d*_*c*,*h*_ to non-adjacent labels. Defining the index set of all samples as *n ∈* {1, 2, …, ∑_*k*_ *N*^(*k*)^}, the centroid for label *c* is as follows.

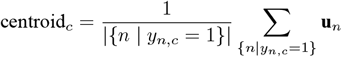

Here, **u**_*n*_ denotes the embedding vector for sample *n*, and *y*_*n*,*c*_ is the (*n, c*)-th element of the matrix **Y**, which consolidates the label information for all samples. The set {*n | y*_*n*,*c*_ = 1} represents the indices of all samples belonging to label *c*, and centroid_*c*_ is the mean embedding vector for label *c*. Using these centroid vectors, the distances *d*_*c*,*c*+1_ and *d*_*c*,*h*_ are defined as follows.

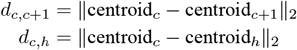

The loss function *L*_*T*_ ensures that the order of samples is preserved in the low-dimensional space by ensuring that *d*_*c*,*c*+1_ is smaller than *d*_*c*,*h*_. A penalty is applied if this condition is violated. Based on the above formulation, the final objective function for the proposed method is as follows.

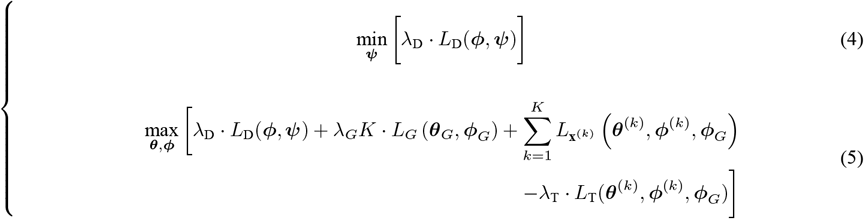

Here, *λ*_*D*_, *λ*_*G*_, and *λ*_*T*_ (*>* 0) are hyperparameters that control the relative importance of adversarial learning, graph-based feature embedding, and the triplet loss term, respectively. Using this objective function, we estimate the parameters via the Adam optimizer [14]. The optimization process is performed in two stages: first, we estimate the parameters related to adversarial learning, followed by estimating parameters associated with the data, graph, and triplet loss terms.

## A Numerical Experiments

In this section, we describe numerical experiments designed to analyze the characteristics of the proposed method. Specifically, we evaluate its effectiveness by generating datasets under multiple conditions and assessing the results of dimensionality reduction and clustering.

### A.1 Data Generation

This subsection explains the procedure for generating synthetic multi-omics datasets used in the numerical experiments. The experiments involve the generation of pseudo scRNA-seq and scATAC-seq data under various conditions, and the performance of the proposed method is evaluated based on different scenarios, which are constructed by combining multiple conditions.

1. **Number of Samples**: The number of samples is set to 200 and 500 to examine the effect of dataset size on method performance. This condition simulates both small and large cell populations and facilitates the assessment of the impact of dataset size.
2. **Number of Features**: The number of features is set to 200 and 500 to evaluate the effect of data dimensionality on performance.
3. **Number of Labels**: The number of label types is set to 5 and 10 to investigate how data diversity influences this method, and the impact of increasing label categories on performance is examined.
4. **Number of Label-Related Features**: The number of features associated with labels is set to 10, 20, and 30 to assess how well the method captures meaningful variations in zero-inflated data. This term refers to datasets commonly observed in scRNA-seq experiments, where most observed values are zero [15].
5. **Label Balance**: The label distribution is set to either “balanced” or “imbalanced” to evaluate the impact of class distribution skewness. In balanced condition, all labels contain the same number of samples. For instance, if a dataset consists of 200 samples and 5 label categories, each label contains 40 samples under balanced condition. If any label contains >40 samples, the dataset is considered imbalanced.
6. **Label Order**: Labels are classified into “unordered” and “ordered” conditions to examine the influence of cell differentiation and expression patterns. In the ordered condition, the dataset is generated such that labels in sequential order have similar values, mimicking pseudotime in cell development.
7. **Dataset Overlap**: The overlap between scRNA-seq and scATAC-seq samples is set to either “100% match” or “50% match” to evaluate how the correspondence between omics datasets affects performance. By adjusting the degree of overlap between scRNA-seq and scATAC-seq datasets, we assess the impact of complementary relationships between omics data.

The details of the data generation process based on the above conditions are described below.

First, we elaborate on the labels assigned to each sample. In Condition 3, the number of labels is set to either 5 or 10, reflecting the diversity of labels within the dataset. When the number of labels is small, distinguishing between classes is relatively straightforward. However, as the number of labels increases, capturing the differences between classes becomes more challenging. This setting allows us to evaluate how different label configurations influence the results.

Then, we describe the label balance. In Condition 5, the label distribution is set to either “balanced” or “imbalanced.” In the balanced setting, each class is assigned an equal number of samples, making label differences more apparent. Conversely, in the imbalanced setting, different numbers of samples are assigned to each label, simulating the class imbalance commonly observed in real-world biological data. The specific sample distributions for the cases with 5 and 10 labels are as follows.

For 5 labels:

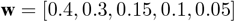

For 10 labels:

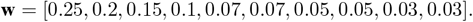

These imbalanced weight distributions generate datasets with significantly different class sizes, enabling us to assess how class imbalance affects the results. Imbalanced datasets can lead to issues such as underestimation and overestimation of minority and majority classes, respectively, making it crucial to evaluating the robustness of the proposed method.

Based on Conditions 1 through 6, we generate synthetic scRNA-seq and scATAC-seq datasets as follows. First, the number of columns that exhibit label-dependent characteristics is restricted to the values specified in Condition 4. This is because real scRNA-seq and scATAC-seq data are often zero-inflated, meaning that only a small fraction of variables contain meaningful values. To reflect this characteristic, we set the number of label-related features to 10, 20, and 30 columns, as specified in Condition 4.

To generate data according to order labels, we divide the dataset’s columns into three categories based on label types and relationships:

- Shared gene set (shared_gene_set): Genes common between adjacent labels
- New gene set (new_gene_set): Genes that are specifically expressed in new labels
- Independent gene set: Genes that differ across all labels

This ensures that the generated data follows the specified order relationships. The corresponding values are generated using a negative binomial distribution NB(*n* = 2, *p* = 1). The entire set of genes is denoted as gene_sets.

The set gene_sets represents the entire collection of columns specified in Condition 4. This set is partitioned into *C* subsets as follows:

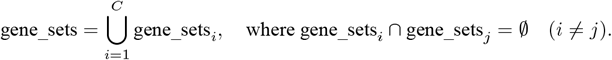

Each subset gene_sets_*i*_ is defined as:

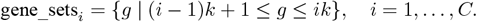

Here, *k* is obtained by taking the integer part of *P* /*C*, with any remainder discarded. Using these definitions, the shared and new gene sets for each label are defined as follows:

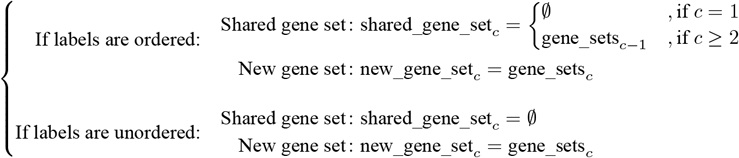

Additionally, to introduce correlations between scRNA-seq and scATAC-seq data, we applied a transformation to adjust the values of scRNA-seq data based on the corresponding scATAC-seq data. Specifically, we randomly selected 70% of columns from the scATAC-seq dataset and created edges linking them to their corresponding columns in the scRNA-seq dataset. Each edge was assigned a random weight weight *~*U(0.1, 1) to represent the interaction strength. Using this graph structure, the scRNA-seq data were updated according to the following equation:

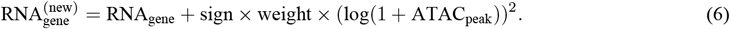

Here, gene and peak denote randomly selected, related columns, where RNA_gene_ represents a specific column in the scRNA-seq dataset, and ATAC_peak_ represents its corresponding column in the scATAC-seq dataset. The term sign denotes the interaction’s direction, while weight *~* U(0.1, 1) is a random parameter representing the interaction strength.

In contrast, for columns that are unspecified in Condition 4 (independent gene sets across labels), the following procedure is applied to generate zero-inflated sparse data. Specifically, for the remaining columns after excluding label-related features specified in Condition 4, 50% of values are set to zero and the remaining 50% are assigned random values. This process results in zero-inflated sparse data, mimicking the characteristics of real gene expression data. Additionally, standard normally distributed noise is added to the generated data.

Finally, we describe the overlap between scRNA-seq and scATAC-seq data in more detail. In Condition 7, the overlap between datasets is set to either “100%” or “50%.” After generating both datasets, the degree of overlap is expressed by modifying sample names. Specifically, when the overlap rate is 100%, the scRNA-seq and scATAC-seq datasets are completely matched, meaning that all samples share identical names. In contrast, when the overlap rate is 50%, half of the samples have the same names in both datasets, whereas the remaining 50% have different names. This operation enables the evaluation of the impact the degree of overlap between scRNA-seq and scATAC-seq datasets on data analysis.

The following summarizes the data generation process described above.

I. **Label Assignment**:
  - Assign cells to labels based on the total number of cells and the number of labels.
    - If label balance is “balanced,” distribute cells equally among all labels.
    - If label balance is “imbalanced,” allocate cells according to predefined proportions.
  - Determine the label structure based on whether labels are “unordered” or “ordered.”
II. **Gene Set Construction**:
  - Define gene sets corresponding to each label based on the number of label-related variables.
    - If label ordering is “ordered,” create shared gene sets between adjacent labels.
III. **Data Generation**:
  - Generate scRNA-seq and scATAC-seq data from a negative binomial distribution for each label.
  - Assign gene expression levels and peak intensities to cells based on gene sets.
    - If dataset overlap is “100% matched,” ensure complete alignment of labels between scRNA-seq and scATAC-seq data.
    - If the dataset overlap is “50% matched,” align only a subset of labels between scRNA-seq and scATAC-seq data.
IV. **Noise Addition**:
  - Add random noise to scRNA-seq and scATAC-seq data.

### A.2 Methods Applied

In this study, we evaluated the performance of the proposed and comparative methods using the following six approaches.

- **GLUE**: GLUE is a method of diagonal integration that has the essential ability to incorporate prior knowledge of feature relationships into the model via a knowledge graph. This allows it to preserve relationships between samples more effectively than other diagonal integration methods [12].
- **CGLUE**: CGLUE is a proposed method that integrates different omics datasets using a model conditioned on labels during embedding by incorporating label information in addition to inter-omics interactions (e.g., between scRNA-seq and scATAC-seq). This produces embeddings that preserve both data structure and the associated label information.
- **CGLUE-SOE**: CGLUE-SOE is an extension of CGLUE that incorporates an additional loss term to enforce order preservation. Additionally, this method adjusts embeddings to align with order labels via a loss function.
- **GLUE-SOE**: GLUE-SOE is a variant of GLUE that adds an order-preserving loss term post-embedding to adjust and preserve the order structure.
- **CCA**: Canonical Correlation Analysis (CCA) is a classical dimensionality reduction method that maximizes linear correlations between omics datasets that is applicable only when the two datasets have fully matched samples. While the CCA does not support diagonal integration or utilize label information, it performs dimensionality reduction based on linear correlations, focusing on shared variations across datasets.
- **UMAP**: Uniform manifold approximation and projection (UMAP) is a nonlinear dimensionality reduction technique that is applied to either a simple concatenation of two datasets with fully matched samples or to a single dataset independently. While UMAP effectively captures local structures and is well-suited for visualization, it does not consider diagonal integration, label information, or order information, performing only basic dimensionality reduction.

The above methods were applied to perform multi-omics data integration and dimensionality reduction. Each method possesses distinct characteristics, which are summarized in Table 3.

**Table 3:** Characteristics of Each Method.

| Method | Embedding | Integration | Diagonal Integration | Label Information | Order Preservation |
| --- | --- | --- | --- | --- | --- |
| CGLUE | ✓ | ✓ | ✓ | ✓ |  |
| GLUE-SOE | ✓ | ✓ | ✓ |  | ✓ |
| CGLUE-SOE | ✓ | ✓ | ✓ | ✓ | ✓ |
| GLUE | ✓ | ✓ | ✓ |  |  |
| CCA | ✓ | ✓ |  |  |  |
| UMAP | ✓ |  |  |  |  |

Table 3 illustrates the specific characteristics of each method. First, all methods perform some form of dimensionality reduction through embedding. However, excluding UMAP, all other methods enable data integration. Diagonal integration is supported by CGLUE, GLUE-SOE, CGLUE-SOE, and GLUE, but not by CCA and UMAP. Methods that support diagonal integration can effectively combine datasets obtained from different sources, allowing for a more comprehensive utilization of diverse data. Regarding the use of label information, only CGLUE and CGLUE-SOE incorporate label information during embedding, allowing them to preserve both data structure and label-based relationships.

For order preservation, only CGLUE-SOE and GLUE-SOE explicitly handle order information, making them suitable for integrating data that reflect cellular differentiation processes or temporal sequences. Other methods do not account for order information, meaning they cannot capture temporal variations or developmental trajectories during data integration.

### A.3 Evaluation Metrics

In this study, we employed two metrics: the ARI to evaluate clustering results and Correlation to measure the agreement between the assigned order and ground truth order labels. The definitions of these metrics are provided below.

First, we describe the ARI, which quantifies the agreement between clustering results and ground truth class labels, adjusted to account for the balance between correct classifications and misclassifications. ARI is defined as follows:

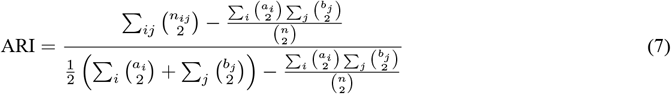

where *n*_*ij*_ represents the number of elements common to cluster *i* and class *j, a*_*i*_ is the number of elements in cluster *i, b*_*j*_ is the number of elements in class *j*, and *n* is the total number of elements. ARI ranges from 1 to 1, with values closer to 1 indicating a stronger agreement between clustering results and true class labels.

In contrast, correlation measures the consistency between the assigned order from clustering and ground truth order labels. Herein, Spearman’s rank correlation coefficient (Correlation) was used and is defined as follows.

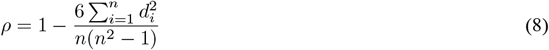

where *d*_*i*_ denotes the difference between the assigned rank and true rank label for element *i*, and *n* represents the total number of elements. The coefficient *ρ* ranges from 1 to 1, with values closer to 1 indicating a stronger agreement between the assigned ranks and ground truth order labels.

These evaluation metrics are used to assess how well the clustering results align with the ground truth labels and their inherent order structure.

### A.4 Evaluation Procedure

In this study, we evaluate the integration performance of scRNA-seq and scATAC-seq data using the six embedding methods described in Section A.2.

To assess the integration performance of each method, we follow the evaluation procedure of [12]. First, scRNA-seq and scATAC-seq data are reduced to a 50-dimensional latent space. Using this embedding, we conduct the following two evaluations. First, *k*-means clustering is performed in the 50-dimensional latent space obtained from each method. This process is repeated 10 times, and the agreement between the assigned cluster labels and ground truth cell labels is measured using ARI.

Second, the 50-dimensional embeddings are further reduced to a two-dimensional space using UMAP. Then, the principal trajectory is inferred in this two-dimensional space using Slingshot [2], and cells are assigned an order. The agreement between the assigned order and ground truth order labels is then evaluated using Spearman’s Correlation.

Thus, the performance of each multi-omics data integration method is assessed in terms of clustering accuracy and order preservation.

Additionally, we primarily evaluate CGLUE-SOE, GLUE-SOE, CGLUE, and GLUE as the main methods. However, UMAP and CCA are also applied as reference methods. These reference methods have limitations that prevent them from being applied to the entire set of scRNA-seq and scATAC-seq samples. Therefore, they are only evaluated under specific conditions. In particular, when the scRNA-seq and scATAC-seq datasets do not completely overlap or when only 50% of samples are shared, the number of usable samples becomes restricted. Therefore, UMAP and CCA are evaluated based on the following approaches, which apply these methods only to the shared data portion.

- **UMAP (only RNA)** applies UMAP only to the scRNA-seq data, performing dimensionality reduction based solely on gene expression levels. Thus, regardless of the overlap rate, only the cells from the scRNA-seq dataset are included in the evaluation.
- **UMAP (only ATAC)** applies UMAP only to the scATAC-seq data, performing dimensionality reduction based solely on chromatin accessibility levels. Similarly, regardless of the overlap rate, only cells from the scATAC-seq dataset are included in the evaluation.
- **UMAP (only intersection)** applies UMAP to the intersection of the scRNA-seq and scATAC-seq datasets. Since this method is limited to overlapping samples when datasets have low overlap, only cells shared between the two datasets are included in the evaluation.
- **CCA** is applied simultaneously to both scRNA-seq and scATAC-seq datasets to maximize the correlation between them. Similar to UMAP (only intersection), only the overlapping samples between the two datasets are included in the evaluation.

Since UMAP and CCA use only one modality or the shared portion of the datasets, their results do not reflect the overall integration performance. Thus, their results should be considered for reference only, rather than to provide direct comparisons with the primary methods.

### A.5 Scenario Overview

**Table 4:** Numerical Experiment Scenarios 1 to 4.

| Element | Scenario 1 | Scenario 2 | Scenario 3 | Scenario 4 |
| --- | --- | --- | --- | --- |
| Number of Samples | 200 rows | 200 rows | 200 rows | 200 rows |
| Number of Variables | 200 columns | 200 columns | 200 columns | 200 columns |
| Number of Labels | 5 | 5 | 5 | 5 |
| Number of Label-Related Variables | 10 columns | 10 columns | 15 columns | 15 columns |
| Label Distribution | Balanced | Imbalanced | Balanced | Imbalanced |
| Label Ordering | Ordered | Ordered | Ordered | Ordered |
| Data Overlap | 100% overlap | 100% overlap | 100% overlap | 100% overlap |

**Table 5:** Numerical Experiment Scenarios 5 to 8.

| Element | Scenario 5 | Scenario 6 | Scenario 7 | Scenario 8 |
| --- | --- | --- | --- | --- |
| Number of Samples | 200 rows | 200 rows | 200 rows | 200 rows |
| Number of Variables | 200 columns | 200 columns | 200 columns | 200 columns |
| Number of Labels | 5 | 5 | 5 | 5 |
| Number of Label-Related Variables | 20 columns | 20 columns | 10 columns | 10 columns |
| Label Distribution | Balanced | Imbalanced | Balanced | Imbalanced |
| Label Ordering | Ordered | Ordered | Ordered | Ordered |
| Data Overlap | 100% overlap | 100% overlap | 50% overlap | 50% overlap |

**Table 6:** Numerical Experiment Scenarios 9 to 12.

| Element | Scenario 9 | Scenario 10 | Scenario 11 | Scenario 12 |
| --- | --- | --- | --- | --- |
| Number of Samples | 200 rows | 200 rows | 200 rows | 200 rows |
| Number of Variables | 200 columns | 200 columns | 200 columns | 200 columns |
| Number of Labels | 5 | 5 | 5 | 5 |
| Number of Label-Related Variables | 15 columns | 15 columns | 20 columns | 20 columns |
| Label Distribution | Balanced | Imbalanced | Balanced | Imbalanced |
| Label Ordering | Ordered | Ordered | Ordered | Ordered |
| Data Overlap | 50% overlap | 50% overlap | 50% overlap | 50% overlap |

**Table 7:** Numerical Experiment Scenarios 13 to 16.

| Element | Scenario 13 | Scenario 14 | Scenario 15 | Scenario 16 |
| --- | --- | --- | --- | --- |
| Number of Samples | 500 rows | 500 rows | 500 rows | 500 rows |
| Number of Variables | 500 columns | 500 columns | 500 columns | 500 columns |
| Number of Labels | 5 | 5 | 5 | 5 |
| Number of Label-Related Variables | 10 columns | 10 columns | 15 columns | 15 columns |
| Label Distribution | Balanced | Imbalanced | Balanced | Imbalanced |
| Label Ordering | Ordered | Ordered | Ordered | Ordered |
| Data Overlap | 100% overlap | 100% overlap | 100% overlap | 100% overlap |

**Table 8:** Numerical Experiment Scenarios 17 to 20.

| Element | Scenario 17 | Scenario 18 | Scenario 19 | Scenario 20 |
| --- | --- | --- | --- | --- |
| Number of Samples | 500 rows | 500 rows | 500 rows | 500 rows |
| Number of Variables | 500 columns | 500 columns | 500 columns | 500 columns |
| Number of Labels | 5 | 5 | 5 | 5 |
| Number of Label-Related Variables | 20 columns | 20 columns | 10 columns | 10 columns |
| Label Distribution | Balanced | Imbalanced | Balanced | Imbalanced |
| Label Ordering | Ordered | Ordered | Ordered | Ordered |
| Data Overlap | 100% overlap | 100% overlap | 50% overlap | 50% overlap |

**Table 9:** Numerical Experiment Scenarios 21 to 24.

| Element | Scenario 21 | Scenario 22 | Scenario 23 | Scenario 24 |
| --- | --- | --- | --- | --- |
| Number of Samples | 500 rows | 500 rows | 500 rows | 500 rows |
| Number of Variables | 500 columns | 500 columns | 500 columns | 500 columns |
| Number of Labels | 5 | 5 | 5 | 5 |
| Number of Label-Related Variables | 15 columns | 15 columns | 20 columns | 20 columns |
| Label Distribution | Balanced | Imbalanced | Balanced | Imbalanced |
| Label Ordering | Ordered | Ordered | Ordered | Ordered |
| Data Overlap | 50% overlap | 50% overlap | 50% overlap | 50% overlap |

### A.6 Results

**Table 10:** Results of numerical experiments for Scenarios 1 and 2.

|  | Scenario 1 |  |  |  | Scenario 2 |  |  |  |
| --- | --- | --- | --- | --- | --- | --- | --- | --- |
|  | Correlation |  | ARI |  | Correlation |  | ARI |  |
|  | Mean | Sd | Mean | Sd | Mean | Sd | Mean | Sd |
| CGLUE-SOE | **0.861** | 0.029 | 0.387 | 0.018 | **0.743** | 0.060 | 0.291 | 0.038 |
| GLUE-SOE | 0.824 | 0.023 | 0.339 | 0.015 | 0.632 | 0.123 | 0.255 | 0.027 |
| CGLUE | 0.859 | 0.024 | **0.389** | 0.021 | 0.740 | 0.078 | **0.299** | 0.034 |
| GLUE | 0.807 | 0.020 | 0.339 | 0.027 | 0.659 | 0.094 | 0.249 | 0.027 |
| UMAP (only RNA) | 0.095 | 0.047 | 0.027 | 0.011 | 0.045 | 0.103 | 0.002 | 0.003 |
| UMAP (only ATAC) | 0.080 | 0.089 | 0.024 | 0.007 | 0.079 | 0.111 | 0.009 | 0.004 |
| UMAP (only intersection) | 0.138 | 0.103 | 0.012 | 0.006 | 0.110 | 0.076 | 0.029 | 0.005 |
| CCA | 0.351 | 0.000 | 0.109 | 0.003 | 0.560 | 0.000 | 0.178 | 0.007 |

**Table 11:** Results of numerical experiments for Scenarios 3 and 4.

|  | Scenario 3 |  |  |  | Scenario 4 |  |  |  |
| --- | --- | --- | --- | --- | --- | --- | --- | --- |
|  | Correlation |  | ARI |  | Correlation |  | ARI |  |
|  | Mean | Sd | Mean | Sd | Mean | Sd | Mean | Sd |
| CGLUE-SOE | **0.970** | 0.003 | 0.822 | 0.019 | **0.932** | 0.021 | 0.877 | 0.093 |
| GLUE-SOE | 0.960 | 0.004 | 0.768 | 0.017 | 0.883 | 0.028 | 0.895 | 0.040 |
| CGLUE | **0.970** | 0.005 | **0.849** | 0.027 | 0.927 | 0.019 | 0.839 | 0.112 |
| GLUE | 0.949 | 0.003 | 0.776 | 0.031 | 0.882 | 0.040 | **0.900** | 0.024 |
| UMAP (only RNA) | 0.294 | 0.055 | 0.103 | 0.006 | 0.253 | 0.031 | 0.045 | 0.006 |
| UMAP (only ATAC) | 0.032 | 0.157 | 0.011 | 0.005 | 0.168 | 0.082 | 0.030 | 0.010 |
| UMAP (only intersection) | 0.105 | 0.128 | 0.097 | 0.012 | 0.017 | 0.107 | 0.026 | 0.007 |
| CCA | 0.399 | 0.000 | 0.130 | 0.006 | 0.858 | 0.000 | 0.020 | 0.002 |

**Table 12:**
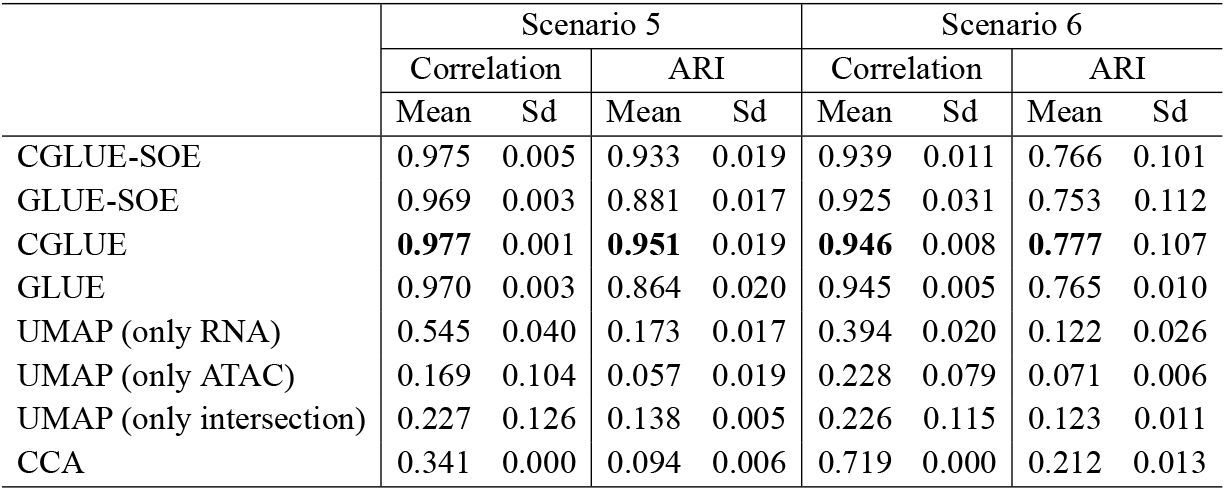
Results of Numerical Experiment Scenarios 5 and 6.

|  | Scenario 5 |  |  |  | Scenario 6 |  |  |  |
| --- | --- | --- | --- | --- | --- | --- | --- | --- |
|  | Correlation |  | ARI |  | Correlation |  | ARI |  |
|  | Mean | Sd | Mean | Sd | Mean | Sd | Mean | Sd |
| CGLUE-SOE | 0.975 | 0.005 | 0.933 | 0.019 | 0.939 | 0.011 | 0.766 | 0.101 |
| GLUE-SOE | 0.969 | 0.003 | 0.881 | 0.017 | 0.925 | 0.031 | 0.753 | 0.112 |
| CGLUE | **0.977** | 0.001 | **0.951** | 0.019 | **0.946** | 0.008 | **0.777** | 0.107 |
| GLUE | 0.970 | 0.003 | 0.864 | 0.020 | 0.945 | 0.005 | 0.765 | 0.010 |
| UMAP (only RNA) | 0.545 | 0.040 | 0.173 | 0.017 | 0.394 | 0.020 | 0.122 | 0.026 |
| UMAP (only ATAC) | 0.169 | 0.104 | 0.057 | 0.019 | 0.228 | 0.079 | 0.071 | 0.006 |
| UMAP (only intersection) | 0.227 | 0.126 | 0.138 | 0.005 | 0.226 | 0.115 | 0.123 | 0.011 |
| CCA | 0.341 | 0.000 | 0.094 | 0.006 | 0.719 | 0.000 | 0.212 | 0.013 |

**Table 13:** Results of Numerical Experiment Scenarios 7 and 8.

|  | Scenario 7 |  |  |  | Scenario 8 |  |  |  |
| --- | --- | --- | --- | --- | --- | --- | --- | --- |
|  | Correlation |  | ARI |  | Correlation |  | ARI |  |
|  | Mean | Sd | Mean | Sd | Mean | Sd | Mean | Sd |
| CGLUE-SOE | 0.849 | 0.038 | **0.384** | 0.027 | **0.837** | 0.025 | **0.346** | 0.031 |
| GLUE-SOE | 0.795 | 0.037 | 0.328 | 0.023 | 0.779 | 0.031 | 0.261 | 0.021 |
| CGLUE | **0.851** | 0.036 | 0.381 | 0.019 | 0.831 | 0.020 | 0.329 | 0.025 |
| GLUE | 0.763 | 0.050 | 0.321 | 0.023 | 0.765 | 0.042 | 0.280 | 0.023 |
| UMAP (only RNA) | 0.050 | 0.076 | 0.003 | 0.005 | 0.004 | 0.070 | 0.015 | 0.005 |
| UMAP (only ATAC) | 0.113 | 0.103 | 0.031 | 0.006 | 0.115 | 0.054 | 0.006 | 0.004 |
| UMAP (only intersection) | -0.072 | 0.098 | 0.005 | 0.010 | 0.099 | 0.061 | 0.009 | 0.008 |
| CCA | 0.371 | 0.000 | 0.100 | 0.017 | 0.647 | 0.000 | 0.193 | 0.033 |

**Table 14:** Results of Numerical Experiment: Scenarios 9 and 10.

|  | Scenario 9 |  |  |  | Scenario 10 |  |  |  |
| --- | --- | --- | --- | --- | --- | --- | --- | --- |
|  | Correlation |  | ARI |  | Correlation |  | ARI |  |
|  | Mean | Sd | Mean | Sd | Mean | Sd | Mean | Sd |
| CGLUE-SOE | **0.974** | 0.003 | **0.869** | 0.030 | **0.924** | 0.020 | 0.906 | 0.023 |
| GLUE-SOE | 0.956 | 0.005 | 0.772 | 0.014 | 0.898 | 0.023 | 0.875 | 0.036 |
| CGLUE | 0.973 | 0.002 | 0.864 | 0.033 | 0.911 | 0.026 | **0.912** | 0.019 |
| GLUE | 0.956 | 0.007 | 0.789 | 0.043 | 0.884 | 0.030 | 0.861 | 0.024 |
| UMAP (only RNA) | 0.172 | 0.064 | 0.116 | 0.015 | 0.261 | 0.074 | 0.056 | 0.007 |
| UMAP (only ATAC) | 0.070 | 0.110 | 0.017 | 0.008 | 0.240 | 0.068 | 0.054 | 0.013 |
| UMAP (only intersection) | 0.023 | 0.110 | 0.024 | 0.012 | 0.140 | 0.070 | 0.036 | 0.016 |
| CCA | 0.426 | 0.000 | 0.109 | 0.013 | 0.762 | 0.000 | 0.385 | 0.080 |

**Table 15:** Results of Numerical Experiment: Scenarios 11 and 12.

|  | Scenario 11 |  |  |  | Scenario 12 |  |  |  |
| --- | --- | --- | --- | --- | --- | --- | --- | --- |
|  | Correlation |  | ARI |  | Correlation |  | ARI |  |
|  | Mean | Sd | Mean | Sd | Mean | Sd | Mean | Sd |
| CGLUE-SOE | **0.976** | 0.002 | 0.905 | 0.045 | **0.949** | 0.001 | 0.958 | 0.012 |
| GLUE-SOE | 0.972 | 0.004 | 0.841 | 0.070 | 0.942 | 0.005 | 0.895 | 0.086 |
| CGLUE | 0.926 | 0.105 | **0.920** | 0.009 | 0.946 | 0.004 | **0.972** | 0.009 |
| GLUE | 0.970 | 0.004 | 0.885 | 0.024 | 0.945 | 0.003 | 0.842 | 0.119 |
| UMAP (only RNA) | 0.548 | 0.057 | 0.170 | 0.010 | 0.415 | 0.081 | 0.163 | 0.049 |
| UMAP (only ATAC) | 0.122 | 0.093 | 0.067 | 0.010 | 0.210 | 0.138 | 0.099 | 0.021 |
| UMAP (only intersection) | 0.221 | 0.079 | 0.078 | 0.015 | 0.089 | 0.060 | 0.031 | 0.006 |
| CCA | 0.544 | 0.000 | 0.379 | 0.067 | 0.817 | 0.000 | 0.488 | 0.074 |

**Table 16:** Results of Numerical Experiment: Scenarios 13 and 14.

|  | Scenario 13 |  |  |  | Scenario 14 |  |  |  |
| --- | --- | --- | --- | --- | --- | --- | --- | --- |
|  | Correlation |  | ARI |  | Correlation |  | ARI |  |
|  | Mean | Sd | Mean | Sd | Mean | Sd | Mean | Sd |
| CGLUE SOE | **0.918** | 0.011 | **0.378** | 0.018 | **0.857** | 0.014 | **0.318** | 0.021 |
| GLUE SOE | 0.833 | 0.009 | 0.266 | 0.012 | 0.680 | 0.029 | 0.170 | 0.008 |
| CGLUE | 0.914 | 0.012 | 0.373 | 0.017 | 0.846 | 0.013 | 0.295 | 0.022 |
| GLUE | 0.830 | 0.020 | 0.269 | 0.012 | 0.688 | 0.046 | 0.197 | 0.015 |
| UMAP (only RNA) | 0.047 | 0.050 | 0.001 | 0.002 | 0.007 | 0.050 | 0.012 | 0.004 |
| UMAP (only ATAC) | 0.048 | 0.047 | 0.003 | 0.001 | 0.053 | 0.025 | 0.010 | 0.003 |
| UMAP (only intersection) | 0.046 | 0.043 | 0.004 | 0.002 | 0.010 | 0.047 | 0.008 | 0.004 |
| CCA | 0.720 | 0.000 | 0.101 | 0.075 | 0.642 | 0.000 | 0.118 | 0.014 |

**Table 17:** Results of Numerical Experiment: Scenarios 15 and 16.

|  | Scenario 15 |  |  |  | Scenario 16 |  |  |  |
| --- | --- | --- | --- | --- | --- | --- | --- | --- |
|  | Correlation |  | ARI |  | Correlation |  | ARI |  |
|  | Mean | Sd | Mean | Sd | Mean | Sd | Mean | Sd |
| CGLUE SOE | **0.978** | 0.000 | 0.907 | 0.024 | **0.948** | 0.002 | 0.875 | 0.112 |
| GLUE SOE | 0.965 | 0.001 | 0.758 | 0.004 | 0.928 | 0.003 | 0.653 | 0.080 |
| CGLUE | **0.978** | 0.001 | **0.917** | 0.011 | **0.948** | 0.002 | **0.949** | 0.014 |
| GLUE | 0.965 | 0.003 | 0.737 | 0.023 | 0.929 | 0.003 | 0.698 | 0.094 |
| UMAP (only RNA) | -0.029 | 0.081 | 0.012 | 0.004 | 0.164 | 0.040 | 0.018 | 0.003 |
| UMAP (only ATAC) | 0.085 | 0.076 | 0.016 | 0.008 | 0.098 | 0.053 | 0.021 | 0.006 |
| UMAP (only intersection) | 0.014 | 0.019 | 0.031 | 0.005 | 0.064 | 0.056 | 0.009 | 0.003 |
| CCA | 0.460 | 0.000 | 0.128 | 0.001 | 0.668 | 0.000 | 0.166 | 0.003 |

**Table 18:** Results of Numerical Experiment Scenarios 17 and 18.

|  | Scenario 17 |  |  |  | Scenario 18 |  |  |  |
| --- | --- | --- | --- | --- | --- | --- | --- | --- |
|  | Correlation |  | ARI |  | Correlation |  | ARI |  |
|  | Mean | Sd | Mean | Sd | Mean | Sd | Mean | Sd |
| CGLUE-SOE | **0.978** | 0.001 | 0.927 | 0.023 | **0.951** | 0.000 | 0.983 | 0.005 |
| GLUE-SOE | 0.968 | 0.004 | 0.823 | 0.018 | 0.925 | 0.064 | 0.786 | 0.121 |
| CGLUE | **0.978** | 0.001 | **0.931** | 0.017 | **0.951** | 0.000 | **0.986** | 0.006 |
| GLUE | 0.970 | 0.003 | 0.819 | 0.051 | 0.945 | 0.002 | 0.859 | 0.115 |
| UMAP (only RNA) | 0.189 | 0.056 | 0.052 | 0.006 | 0.192 | 0.034 | 0.034 | 0.004 |
| UMAP (only ATAC) | 0.087 | 0.057 | 0.020 | 0.004 | 0.116 | 0.073 | 0.013 | 0.003 |
| UMAP (only intersection) | 0.029 | 0.069 | 0.048 | 0.005 | 0.109 | 0.065 | 0.013 | 0.002 |
| CCA | 0.467 | 0.000 | 0.096 | 0.000 | 0.723 | 0.000 | 0.203 | 0.000 |

**Table 19:** Results of Numerical Experiment Scenarios 19 and 20.

|  | Scenario 19 |  |  |  | Scenario 20 |  |  |  |
| --- | --- | --- | --- | --- | --- | --- | --- | --- |
|  | Correlation |  | ARI |  | Correlation |  | ARI |  |
|  | Mean | Sd | Mean | Sd | Mean | Sd | Mean | Sd |
| CGLUE-SOE | 0.909 | 0.011 | **0.358** | 0.021 | **0.849** | 0.017 | 0.287 | 0.021 |
| GLUE-SOE | 0.828 | 0.008 | 0.250 | 0.011 | 0.727 | 0.036 | 0.181 | 0.017 |
| CGLUE | **0.914** | 0.007 | 0.354 | 0.021 | 0.844 | 0.011 | **0.294** | 0.012 |
| GLUE | 0.837 | 0.011 | 0.265 | 0.017 | 0.717 | 0.029 | 0.201 | 0.026 |
| UMAP (only RNA) | 0.107 | 0.019 | 0.005 | 0.002 | 0.077 | 0.044 | −0.001 | 0.005 |
| UMAP (only ATAC) | 0.053 | 0.045 | 0.004 | 0.001 | 0.000 | 0.029 | 0.003 | 0.003 |
| UMAP (only intersection) | 0.039 | 0.047 | 0.008 | 0.004 | 0.124 | 0.075 | 0.007 | 0.005 |
| CCA | 0.322 | 0.000 | 0.076 | 0.004 | 0.778 | 0.000 | 0.267 | 0.041 |

**Table 20:** Results of Numerical Experiment Scenario 21 and Scenario 22.

|  | Scenario 21 |  |  |  | Scenario 22 |  |  |  |
| --- | --- | --- | --- | --- | --- | --- | --- | --- |
|  | Correlation |  | ARI |  | Correlation |  | ARI |  |
|  | Mean | Sd | Mean | Sd | Mean | Sd | Mean | Sd |
| CGLUE-SOE | **0.976** | 0.001 | **0.875** | 0.017 | **0.949** | 0.001 | 0.924 | 0.083 |
| GLUE-SOE | 0.958 | 0.002 | 0.693 | 0.028 | 0.926 | 0.004 | 0.618 | 0.007 |
| CGLUE | **0.976** | 0.001 | 0.873 | 0.022 | 0.948 | 0.003 | **0.947** | 0.012 |
| GLUE | 0.957 | 0.004 | 0.679 | 0.030 | 0.929 | 0.004 | 0.653 | 0.053 |
| UMAP (only RNA) | 0.164 | 0.055 | 0.033 | 0.007 | 0.135 | 0.057 | 0.015 | 0.004 |
| UMAP (only ATAC) | 0.112 | 0.031 | 0.012 | 0.002 | 0.117 | 0.046 | 0.022 | 0.005 |
| UMAP (only intersection) | −0.049 | 0.077 | 0.023 | 0.003 | 0.014 | 0.060 | 0.001 | 0.003 |
| CCA | 0.509 | 0.000 | 0.082 | 0.007 | 0.523 | 0.073 | 0.083 | 0.016 |

**Table 21:** Results of Numerical Experiment Scenarios 23 and 24.

|  | Scenario 23 |  |  |  | Scenario 24 |  |  |  |
| --- | --- | --- | --- | --- | --- | --- | --- | --- |
|  | Correlation |  | ARI |  | Correlation |  | ARI |  |
|  | Mean | Sd | Mean | Sd | Mean | Sd | Mean | Sd |
| CGLUE-SOE | 0.978 | 0.000 | 0.935 | 0.012 | **0.951** | 0.000 | **0.991** | 0.005 |
| GLUE-SOE | 0.974 | 0.002 | 0.866 | 0.016 | 0.946 | 0.004 | 0.930 | 0.082 |
| CGLUE | **0.979** | 0.000 | **0.936** | 0.011 | **0.951** | 0.000 | 0.988 | 0.006 |
| GLUE | 0.972 | 0.003 | 0.861 | 0.018 | 0.948 | 0.002 | 0.898 | 0.102 |
| UMAP (only RNA) | 0.306 | 0.047 | 0.072 | 0.006 | 0.203 | 0.085 | 0.028 | 0.009 |
| UMAP (only ATAC) | 0.159 | 0.045 | 0.031 | 0.005 | 0.087 | 0.056 | 0.007 | 0.002 |
| UMAP (only intersection) | −0.009 | 0.033 | 0.010 | 0.004 | 0.092 | 0.084 | 0.032 | 0.004 |
| CCA | 0.469 | 0.000 | 0.106 | 0.006 | 0.598 | 0.000 | 0.106 | 0.014 |

